# Proteinase-Activated Receptor 4 (PAR4) Activation Triggers Cell Membrane Blebbing through RhoA and β-arrestin

**DOI:** 10.1101/746677

**Authors:** Christina MG Vanderboor, Pierre E Thibeault, Kevin CJ Nixon, Robert Gros, Jamie Kramer, Rithwik Ramachandran

**Affiliations:** Department of Physiology and Pharmacology, Schulich School of Medicine and Dentistry, University of Western Ontario, London, Ontario, Canada

## Abstract

Proteinase-Activated Receptors (PARs) are a four-member family of G-protein coupled receptors that are activated via proteolysis. PAR4 is a member of this family that is cleaved and activated by serine proteinases such as thrombin, trypsin and cathepsin-G. PAR4 is expressed in a variety of tissues and cell types including platelets, vascular smooth muscle cells and neuronal cells. In studying PAR4 signalling and trafficking, we observed dynamic changes in the cell membrane with spherical membrane protrusions that resemble plasma membrane blebbing. Since non-apoptotic membrane blebbing is now recognized as an important regulator of cell migration, cancer cell invasion, and vesicular content release we sought to elucidate the signalling pathway downstream of PAR4 activation that leads to such events. Using a combination of pharmacological inhibition and CRISPR/Cas9-mediated gene-editing approaches we establish that PAR4-dependent membrane blebbing occurs independently of the Gα_q/11_ and Gα_i_ signalling pathways and is dependent on signalling via the β-arrestin-1/-2 and RhoA signalling pathways. In order to gain a more comprehensive understanding of β-arrestin-mediated signalling downstream of PAR4 and to guide future studies, we undertook RNA-seq analysis of PAR4 activated genes in control cells and in cells lacking β-arrestin-1/-2. A list of differentially expressed genes was generated followed by Gene Ontology (GO) and enrichment analysis, revealing PAR4 regulation of genes involved in processes including blood coagulation and circulation, cell-cell adhesion, sensory perception and neuron-neuron synaptic transmission-terms that relate back to known functions of PAR4 and are consistent with our finding of membrane blebbing triggered by PAR4 activation. Together these studies provide further mechanistic insight into PAR4 regulation of cellular function.

**Significance Statement:** We find that the thrombin receptor PAR4 triggers cell membrane blebbing in a RhoA- and β-arrestin-dependent manner. In addition to identifying novel cellular responses mediated by PAR4, these data provide further evidence for biased signaling in PAR4 since membrane blebbing was dependent on some, but not all, signaling pathways activated by PAR4. Finally through CRISPR/Cas9-mediated targeting and RNA-seq analysis we catalogue here PAR4-dependent transcription that is dependent on β-arrestin.

## Introduction

Proteinase activated receptors (PARs) are a four-member family of G-protein coupled receptors (GPCRs). PARs are unique among GPCRs, in being activated via proteolytic unmasking of a receptor-activating ‘tethered ligand’, that interacts intramolecularly with the orthosteric ligand binding pocket to trigger signalling (Ramachandran *et al.*, 2012). PAR4, the most recently identified member of this family (Xu *et al.*, 1998), is expressed in a variety of tissues and cell types including the platelets, vascular smooth muscle cells, neuronal cells, and some cancer cells.

Much work has been done to develop PAR1- and PAR4-targeted compounds as anti-platelet agents. Both PAR1 and PAR4 are expressed in human platelets and both of these receptors are activated by the coagulation cascade enzyme thrombin. Importantly though, PAR1 and PAR4 appear to serve different roles in the platelet activation process (Kahn *et al.*, 1999), with PAR1 the high-affinity thrombin receptor playing an initiating role and the lower-affinity thrombin receptor PAR4 serving to consolidate and propagate the clot (Kahn *et al.*, 1999). The PAR1 antagonist voropaxar (zontivity), while highly effective in reducing cardiovascular complications, exhibited significant side effects with an elevated risk for bleeding including in the brain (Morrow *et al.*, 2012), further spurring recent efforts to target PAR4. Recent work with small molecule PAR4 antagonists has supported the idea that PAR4 antagonists are effective in reducing platelet rich thrombus formation in human platelets *ex vivo* (Wilson *et al.*, 2017) and in rodent and non-human primate models *in vivo* (Wong *et al.*, 2017). In non-human primate models, PAR4 blockade was associated with low bleeding liability and had a markedly wider therapeutic window compared to the commonly used antiplatelet agent clopidogrel (Wong *et al.*, 2017).

In keeping with emerging literature for other GPCRs, we now know that activated PAR4 can directly couple to multiple G-protein-signalling pathways including Gα_q/11_ and the Gα_12/13_ pathway (Woulfe, 2005; Kim *et al.*, 2006) but is thought not to engage Gα_i_ dependent signalling pathways (Kim *et al.*, 2006). PAR4 can also recruit and signal through β-arrestins (Li *et al.*, 2011; Ramachandran *et al.*, 2017). In recent work, we identified a C-terminal motif in PAR4 that was critical for PAR4 signalling through the Gα_q/11_ calcium signalling pathway and for recruiting β-arrestin-1/-2 (Ramachandran *et al.*, 2017). The mutant receptor with an 8-amino acid C-terminal deletion (dRS-PAR4) failed to internalize following activation with the PAR4 agonists thrombin or AYPGKF-NH_2_ suggesting a role for β-arrestins in PAR4 trafficking. A pepducin targeting this C-terminal motif was also effective in attenuating PAR4-dependent platelet aggregation and thrombosis *in vivo* (Ramachandran *et al.*, 2017). These recent findings point to the exciting possibility that it might be possible to therapeutically target PAR4 signalling in a pathway-specific manner. The present study was spurred by our observations that PAR4 activation rapidly triggered the formation of dynamic membrane blebs, which were absent in dRS-PAR4 expressing cells.

Non-apoptotic plasma membrane blebbing is now recognized as a feature of various cellular processes including directional cellular migration during development, cancer cell migration and invasion, neuronal cell remodeling, and vesicular content release (Charras, 2008; Charras and Paluch, 2008; Charras *et al.*, 2008). Blebs are formed when the plasma membrane transiently detaches from the underlying actin filaments resulting in intracellular pressure-mediated spherical membrane protrusions (Charras *et al.*, 2008; Tinevez *et al.*, 2009). The reassembly of actin filaments limits the expansion of blebs and actin polymerization while actomyosin contraction drives the retraction of the blebs (Charras *et al.*, 2008). The molecular signals that trigger the formation of membrane blebs are begining to be understood and include signalling from cell surface receptor such as GPCRs and receptor tyrosine kinases (Hagmann *et al.*, 1999; Lawrenson *et al.*, 2002; Godin and Ferguson, 2010; Chen *et al.*, 2012; Laser-Azogui *et al.*, 2014). A role for multiple Rho isoforms has also been described in regulating various aspects of bleb formation and retraction (Pinner and Sahai, 2008; Aoki *et al.*, 2016; Gong *et al.*, 2018). Previous work has described regulation of Rho-signalling by GPCRs in both a G-protein- and β-arrestin-dependent manner (Sah *et al.*, 2000; Barnes *et al.*, 2005; Anthony *et al.*, 2011).

Here we examine in detail the pathways leading from PAR4 to the formation of membrane blebs. We find that inhibition of Gα_q/11_ signalling had no effect on the formation of PAR4 triggered membrane blebs, while blockade of β-arrestin-1/-2- or Rho-dependent signalling significantly reduced blebbing.

## Materials and Methods

### Materials

DMEM, trypsin-EDTA (0.25%), PBS, Penicillin-Streptomycin and sodium pyruvate were purchased from Thermo Fisher (Waltham, MA). Peptide ligands were custom synthesized by Genscript (Piscataway, NJ) at greater than 95% purity. Thrombin was purchased from Calbiochem (Oakville, ON), coelanterazine-h was from Nanolight Technology (Pinetop, AZ). All antibodies (anti-β-arrestin-1/-2, anti-RhoA, anti-actin, anti-rabbit-HRP) used in this study were purchased from Cell Signalling Technologies. YM254890 was from Wako chemicals (Richmond, VA), GSK269962 was from Tocris (Oakville, ON) and all other chemicals were from Sigma-Aldrich (Oakville, ON).

### Cell Culture and transfections

HEK-293 (ATTC) cells were maintained in DMEM (Gibco) with 10% fetal bovine serum (Gibco), 1% penicillin-streptomycin (Gibco) and 1% sodium pyruvate (Gibco). Cells stably expressing PAR4-YFP or PAR4-dRS-YFP were maintained in the above media supplemented with 600 μg/mL of geneticin (Gibco). Cells were transiently transfected using a modified calcium phosphate method (Ferguson and Caron, 2004). Experiments were performed 48 hours post transfection. Primary rat smooth muscle cells were isolated and cultured as previously described (Gros *et al.*, 2006).

### Creation of β-arrestin-1/-2 and RhoA knockout cells

RhoA and β-Arrestin-1/-2 knock out HEK-293 cells (RhoA-KO HEK and β-arrestin-1/-2-KO HEK respectively) were generated using CRISPR/Cas9 mediated gene targeting. Guides targeting the β-arrestins or RhoA were designed using the design tool at http://crispr.mit.edu. Gene specific guides (β-arrestin 1; TGTGGACCACATCGACCTCG, β-arrestin 2; GCGGGACTTCGTAGATCACC and RhoA; CGGTCCGCGAGTCGCAAACT, GAGTCCAGCCTCTTCGCGCC, GACTCGCGGACCGGCGTCCC) were ligated into the PX458 vector (a kind gift from Dr. Feng Zhang, MIT, Addgene plasmid # 48138), verified by direct sequencing and transfected in HEK cells via the calcium phosphate method. 48 hours post transfection, GFP expressing single cells were flow sorted into individual wells of a 96 well (Becton Dickinson FACSAria III). Clonal cells from individual wells were screened by western blotting to identify cell lines which were deficient in β-arrestin 1/2 or RhoA (Supplementary Figure 1).

**Figure 1.**
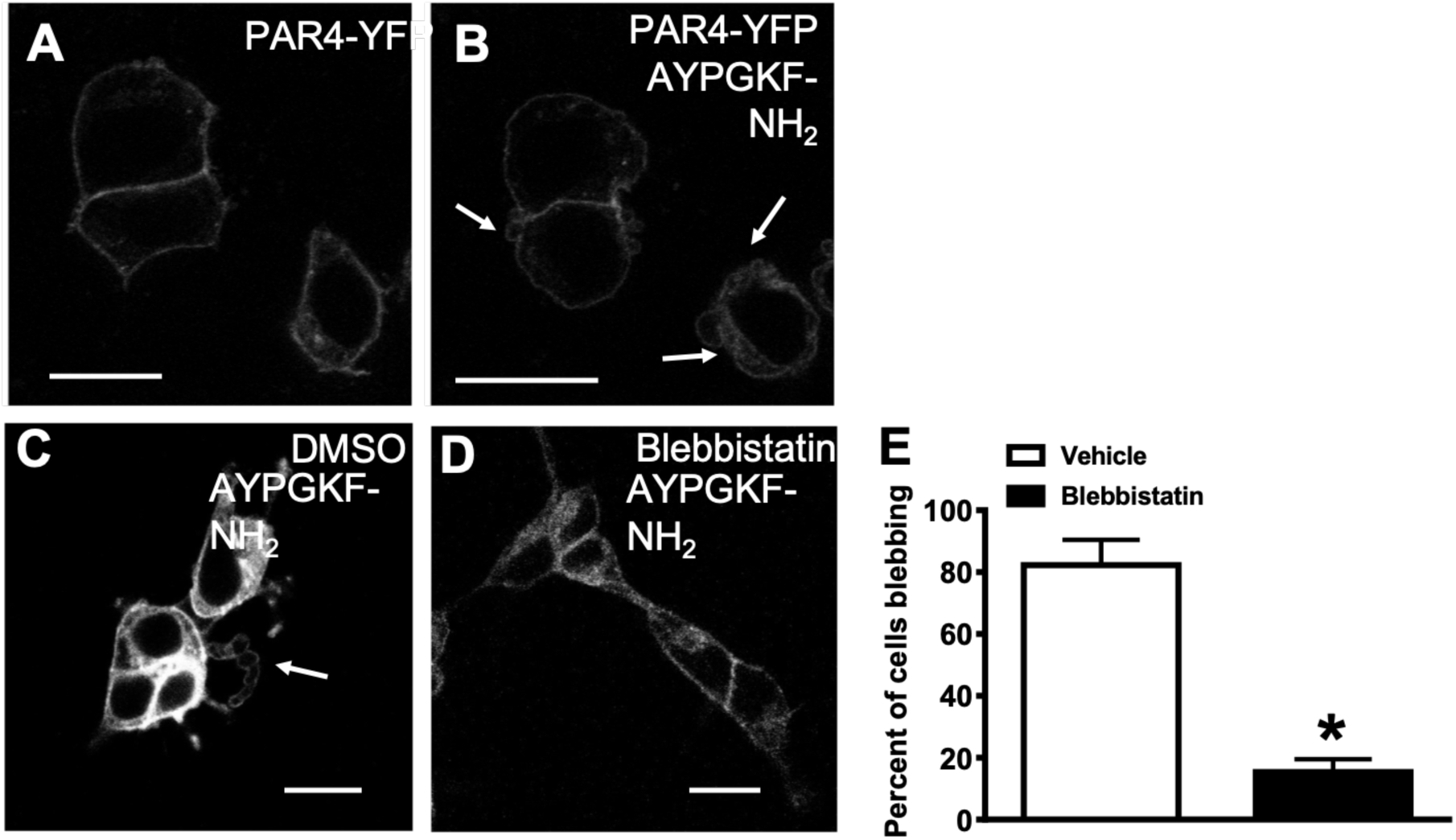
Activation PAR4 mediates a cell shape change response. Representative confocal micrographs, showing HEK-293 cell stably expressing PAR4-YFP (A, B) Cells were treated with 30 μM AYPGKF-NH_2_ for 2 minutes prior to imaging. Size bars are 20 μm, arrows show cell shape change. HEK-293 cells stably expressing PAR4-YFP (C-E) were incubated in either DMSO (C) or 10 μM blebbistatin (B) for 15 minutes prior to a 2-minute treatment with 30 μM AYPGKF-NH_2_ followed by confocal microscopy. Arrows show bleb formation, size bars are 20 μm. (E) graph shows mean +/- SEM, n = 4, asterisk shows significantly different, Mann-Whitney, p<0.05.

### Confocal microscopy

Cells were plated onto glass bottom 35 mm dishes (MatTek, Ashland, MA) and imaged using Zeiss LSM 510 Meta NLO confocal microscope. Yellow fluorescent protein was excited with 514 laser line and visualized with 535-560 filter set. mCherry fluorophore was excited with 543 laser line and visualized with 560-590. Green fluorescent protein was excited with 488 laser line and visualized with 530-560 filter set. Cell shape change experiments were conducted as follows. Cells were treated with vehicle or inhibitor as indicated in the figure legends and incubated at 37°C. Plates were then placed on a heated stage on the microscope and PAR4 was activated with AYPGKF-NH_2_ (30μM) or thrombin (3U/ml). 6-12 images per dish were taken over 10 minutes. Images were analyzed by manually counting cells displaying membrane protrusions or blebs versus cells that did not display any membrane protrusions.

### Bioluminescent Resonance Energy Transfer (BRET) assay for β-arrestin-1/-2 recruitment

Bioluminescent resonance energy transfer was measured between c-terminally YFP tagged PAR4 (Ramachandran *et al.*, 2017) and Renilla Luciferase (Rluc) tagged β-arrestin-1 or β-arrestin-2 (a kind gift from Dr. Michel Bouvier, U. de Montreal), following 20 minutes of receptor activation as described in previous studies (Ramachandran *et al.*, 2009). Briefly, PAR4-YFP (1 µg) and β-arr-1-Rluc and -2-Rluc) (0.1µg) were transiently transfected in cells plated in a six well plate for 24 hours. Cells were re-plated into white 96-well culture plates and cultured for a further 24 hours. Interactions between PAR4 and β-arrestin-1/-2 were detected by measuring the BRET ratio at timed intervals over 20 min following the addition of 5 µM coelenterazine (Nanolight Technology, Pinetop, AZ) on a Mithras fluorescence plate reader (Berthold) in luminescence mode using the appropriate filters.

### RNA-seq PAR4 transcriptome analysis in control and β-arrestin-1/-2 knockout HEK-293 cells

HEK-293 cells or β-arrestin-1/-2-KO HEK cells stably expressing PAR4-YFP were treated with 30μM AYPGKF-NH_2_ for 2 hours. RNA was extracted using an RNeasy mini kit (Qiagen) and analyzed on a bioanalyzer (Agilent, Santa Clara, CA). Sequencing libraries were generated using the ScriptSeq complete kit which incorporated the Ribo-Zero ribosomal RNA (rRNA) removal technology. Sequencing was done on a NextSeq (Illumina) at the London Regional Genomics Centre.

### Sequencing Analysis

Raw sequencing reads were trimmed using Prinseq quality trimming (Schmieder and Edwards, 2011) with a minimum base quality score of 30. Read quality was then assessed using FastQC (http://www.bioinformatics.babraham.ac.uk/projects/fastqc). Trimmed reads were then aligned to the *Homo sapiens* reference genome (CRCh release 38) using the STAR RNA-Seq aligner (Dobin *et al.*, 2013). An average of 28,049,697 and 20,706,493 high quality uniquely aligned reads with a maximum of four mismatches were obtained from HEK-293 cells and β-arrestin-1/-2-KO-HEK cells respectively. (Table 1 and Figure S2 and S3). The number of reads per gene was quantified by inputting reads that uniquely aligned to the reference genome with a maximum of four mismatches into HTSeq-count (Anders *et al.*, 2015) with the - type flag indicating ‘gene’ (Table 1 and Figure S2 and S3). Genes associated with rRNA and non-protein coding genes were removed from the analysis and genes with no reads across all samples in the count tables were removed from analysis leaving 18,615 genes for downstream analysis. Raw gene counts were normalized and differential expression analysis between HEK-293 cells and β-arrestin-1/-2-KO HEK cells with and without the treatment of 30μM AYPGKF-NH_2_ was performed using the R (R: A language and environment for statistical computing. Vienna, Austria: R Foundation for Statistical Computing; 2017) package DESEQ2 (Love *et al.*, 2014). Differentially expressed genes were defined as genes with a fold-change >2 and a Benjamini-Hochberg adjusted p-value <0.05. Overlap between datasets was determined and visualized as a Venn diagram using BioVenn (Zhang *et al.*, 2009). Gene ontology (GO) enrichment analysis was performed on upregulated and downregulated differentially expressed genes with the Panther software (Mi *et al.*, 2017) on http://geneontology.org using the GO-SLIM function.

**Table 1:** Differential PAR4-mediated gene regulation in β-arrestin-1/-2 knock out cells. Gene ontology (GO) biological process terms for genes upregulated (A) and downregulated (B) genes in β-arrestin-1/-2-KO HEK cells transiently expressing PAR4-YFP treated with 30 μM AYPGKF-NH_2_ for 3 hours compared to HEK-293 cells under the same conditions. Top ten significant terms (p<0.05) where the highest enrichment for biological process are presented. Terms related to the membrane blebbing phenotype are in bold and terms related to known functions of PAR4 are italicized.

| Gene Ontology Term | Fold Enrichment |
| --- | --- |
| *Neuron-neuron synaptic transmission (GO:0007270)* | 7.69 |
| Natural killer cell activation (GO:0030101) | 5.45 |
| Fertilization (GO:0009566) | 4.68 |
| Sensory perception of taste (GO:0050909) | 4.11 |
| ***Blood coagulation (GO:0007596)*** | 3.1 |
| ***Cell-cell adhesion (GO:0016337)*** | 2.81 |
| Sensory perception of chemical stimulus (GO:000760) | 2.8 |
| Macrophage activation (GO:0042116) | 2.78 |
| ***Blood circulation (GO:0008015)*** | 2.73 |
| *Sensory perception (GO:0007600)* | 2.63 |

**Table 1**
| Gene Ontology Term | Fold Enrichment |
| --- | --- |
| JNK cascade (GO:0007254) | 7.88 |
| Anatomical structure morphogenesis (GO:0009653) | 3.85 |
| Death (GO:0016265) | 3.06 |
| Cell death (GO:0008219) | 3.06 |
| Apoptotic process (GO:0006915) | 3.01 |
| **Intracellular signal transduction (GO:0035556)** | 2.89 |
| Immune system process (GO:0002376) | 2.77 |
| Cell cycle (GO:0007049) | 2.36 |
| Developmental process (GO:0032502) | 2.22 |
| **Signal transduction (GO:0007165)** | 1.72 |

### Accession numbers

The raw data for RNA-seq is available at the NCBI Gene Expression omnibus (GEO), accession number GSE134713.

### Statistical Analysis

Statistical analysis of data and curve fitting were done with Prism 7 software (GraphPad Software, San Diego, CA). Statistical significance was assessed using the Student’s t-test or Anova with Tukey’s post-test.

## Results

### Activation of PAR4 elicits cell shape changes that are dependent on a C-tail eight-amino acid sequence

Stimulation of plasma membrane localized PAR4-YFP in HEK-293 cells stably expressing PAR4-YFP with the PAR4 specific peptide agonist AYPGKF-NH_2_ resulted in cell shape changes (Fig. 1A, B). PAR4 expressing cells treated with 30 μM AYPGKF-NH_2_, displayed protrusions forming at the plasma membrane, indicated by arrows (Fig. 1B). These structures resemble membrane blebs and began to form around 2 minutes post agonist stimulation and lasted for up to 30 minutes in the presence of 30 μM AYPGKF-NH_2_. In order to further verify that the cytoskeletal changes were indeed membrane blebs, we examined the effect of treating cells with Blebbistatin, a small molecule inhibitor of myosin II ATPase (Cheung *et al.*, 2002; Straight *et al.*, 2003) which functions by locking actin heads in a low actin affinity complex (Kovács *et al.*, 2004) and is reported to inhibit non-apoptotic membrane blebbing. Incubation of cells with blebbistatin significantly reduced the AYPGKF-NH_2_-stimulated membrane bleb response in PAR4-YFP expressing HEK-293 cells (15.37% +/-4.14) (Fig. 1C-E). In contrast, PAR4-YFP expressing cells treated with DMSO vehicle control displayed membrane blebs with a mean of 82.25% +/-8.25. In recent studies we described a mutant PAR4 receptor lacking eight amino acids from the C-tail, dRS-PAR4-YFP (Ramachandran *et al.*, 2017). We observed that in contrast to the wild type receptor, dRS-PAR4-YFP expressing cells display significantly less blebbing in response to 30 μM AYPGKF-NH_2_ treatment (Fig. 2A-C). 82.33% +/-1.2 of HEK-293 cells stably expressing wild type PAR4-YFP displayed membrane blebbing as opposed to 8.2% +/-2.35 dRS-PAR4-YFP expressing cells, indicating that PAR4 triggered membrane blebbing required the activation of signalling pathways that are dependent on the eight-amino acid sequence in the C-tail of PAR4. Previously, we established that dRS-PAR4-YFP does not couple to Gα_q/11_ and is unable to recruit β-arrestins in response to thrombin or AYPGKF-NH_2_ activation (Ramachandran *et al.*, 2017). Since this mutant receptor is also unable to activate blebbing, we hypothesized that PAR4 cell shape changes are Gα_q/11_- and/or β-arrestin-dependent and examined the effect of blocking these pathways on bleb formation.

**Figure 2.**
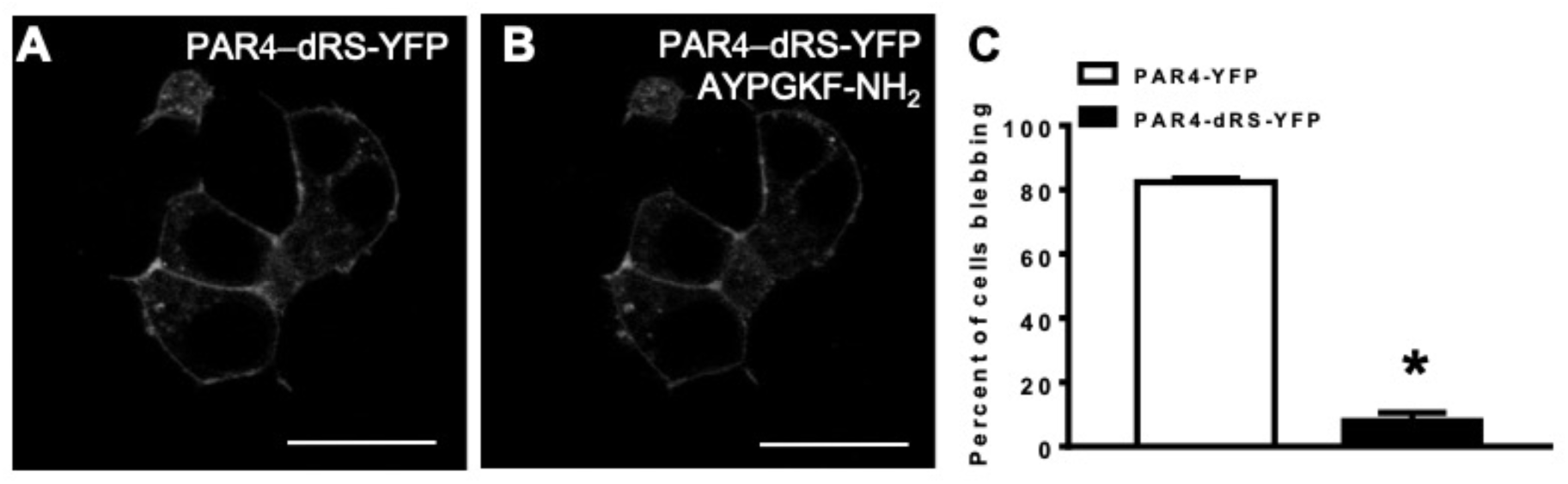
Activation PAR4 mediates a cell shape change response is dependent on an 8 amino acid sequence. HEK-293 transiently expressing dRS-PAR4-YFP (A-C) were treated with 30 μM AYPGKF-NH_2_ for 2 minutes prior to imaging and scored for cell shape change or no change, graph shows mean +/- SEM, size bars are 20 μm, asterisks indicates statistically significant, t-test, n = 3, p<0.005.

### PAR4-mediated cell shape change is Gα_q/11_ and Gα_i_ independent

In order to determine whether PAR4 triggered blebbing is Gα_q/11_-dependent, we treated cells with the potent and selective Gα_q/11_ inhibitor YM-254890 (Taniguchi *et al.*, 2004). It is well established that GPCRs couple to Gα_q/11_ to mobilize calcium and activate protein kinase C (PKC) (Exton, 1996; Wettschureck and Offermanns, 2005). Activated-PKC translocation from the cytosol to the plasma membrane can be observed to monitor this process (Dale *et al.*, 2001; Policha *et al.*, 2006). We employed this assay to visualize the efficacy of YM-254890 in inhibiting Gα_q/11_ signalling through PAR4. Cells were transiently transfected with PAR4-mCherry and PKCβ1-GFP. PAR4 expression was observed at the cell membrane and PKCβ1 expression was evident in the cytoplasm in resting cells. Upon treatment with 30 μM AYPGKF-NH_2,_ PKCβ1-GFP translocated to the membrane (Fig. 3A-C) and blebbing responses were observed, as before. In cells treated with 100 nM YM-254890, PKCβ1-GFP failed to translocate to the membrane following treatment with AYPGKF-NH_2_. 100nM YM-254890 treated cells however maintained their ability to bleb in response to AYPGKF-NH_2_ (Fig. 3D-F). The role of Gα_q/11_ in PAR4-mediated blebbing was further quantified in HEK-293 cells stably expressing PAR4-YFP treated with either DMSO vehicle or YM-254890 prior to stimulation with 30 μM AYPGKF-NH_2_. Blebbing in vehicle treated cells was not significantly different from cells treated with YM-254890, 81.0% +/-4.5 and 77.0% +/-2.5 respectively (Fig. 3G). This data indicates that YM-254890 functionally blocks Gα_q/11_ signalling as indicated by a lack of PKCβ1 translocation but does not block cell shape changes mediated by PAR4 activation.

**Figure 3.**
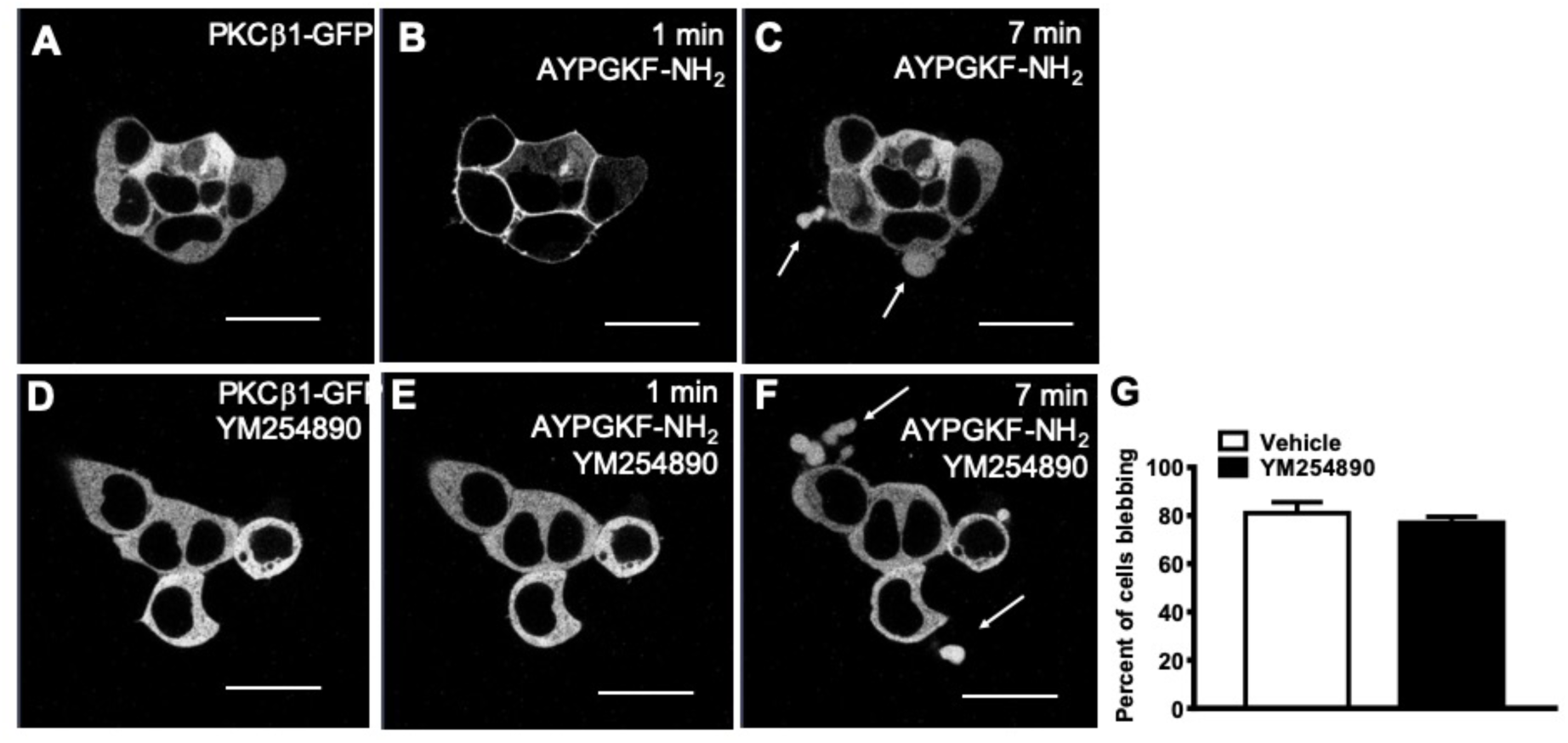
PAR4-mediated cell shape change is Gα_q/11_-independent. HEK-293 cells transiently expressing PAR4-mCherry (not shown) with PKCβ1-GFP, were treated with DMSO followed by stimulation with 30 μM AYPGKF-NH_2_ (A-C) or with 100 nM YM254890 for 20 minutes followed by stimulation with 30 μM AYPGKF-NH_2_ (D-F). Arrows show bleb formation, size bars are 20 μm. Graph shows mean +/- SEM, n = 3, not significantly different, t-test (G).

HEK-293 cells, transiently expressing PAR4-mCherry with PKCβ1-GFP, showed that activation of PAR4-mCherry with AYPGKF-NH_2_ causes a translocation of PKCβ1-GFP from the cytosol to the plasma membrane (Fig. 4A, B). In contrast, HEK-293 cells transiently expressing dRS-PAR4-mCherry with PKCβ1-GFP showed that activation of dRS-PAR4-mCherry does not cause a redistribution of PKCβ1-GFP to the membrane (Fig. 4C, D). Although PKC activation is downstream of Gα_q/11_, since dRS-PAR4 does not activate PKC and does not elicit cell shape changes, we tested PKC for a potential role in mediating PAR4-mediated cell shape changes. HEK-293 cells stably expressing PAR4-YFP were treated with the PKC inhibitor Gö6983 (Gschwendt *et al.*, 1996), prior to stimulation with 30 μM AYPGKF-NH_2_ and visualization by confocal microscopy. There was no significant difference in the number of DMSO vehicle treated cells displaying blebbing when compared to cells treated with Gö6983 in response to AYPGKF-NH_2_, 74.5% +/-4.6 and 72.2% +/-9.2 (Fig. 4G). This data then suggests that neither Gα_q/11_ or PKC facilitate PAR4-mediated cell shape changes.

**Figure 4.**
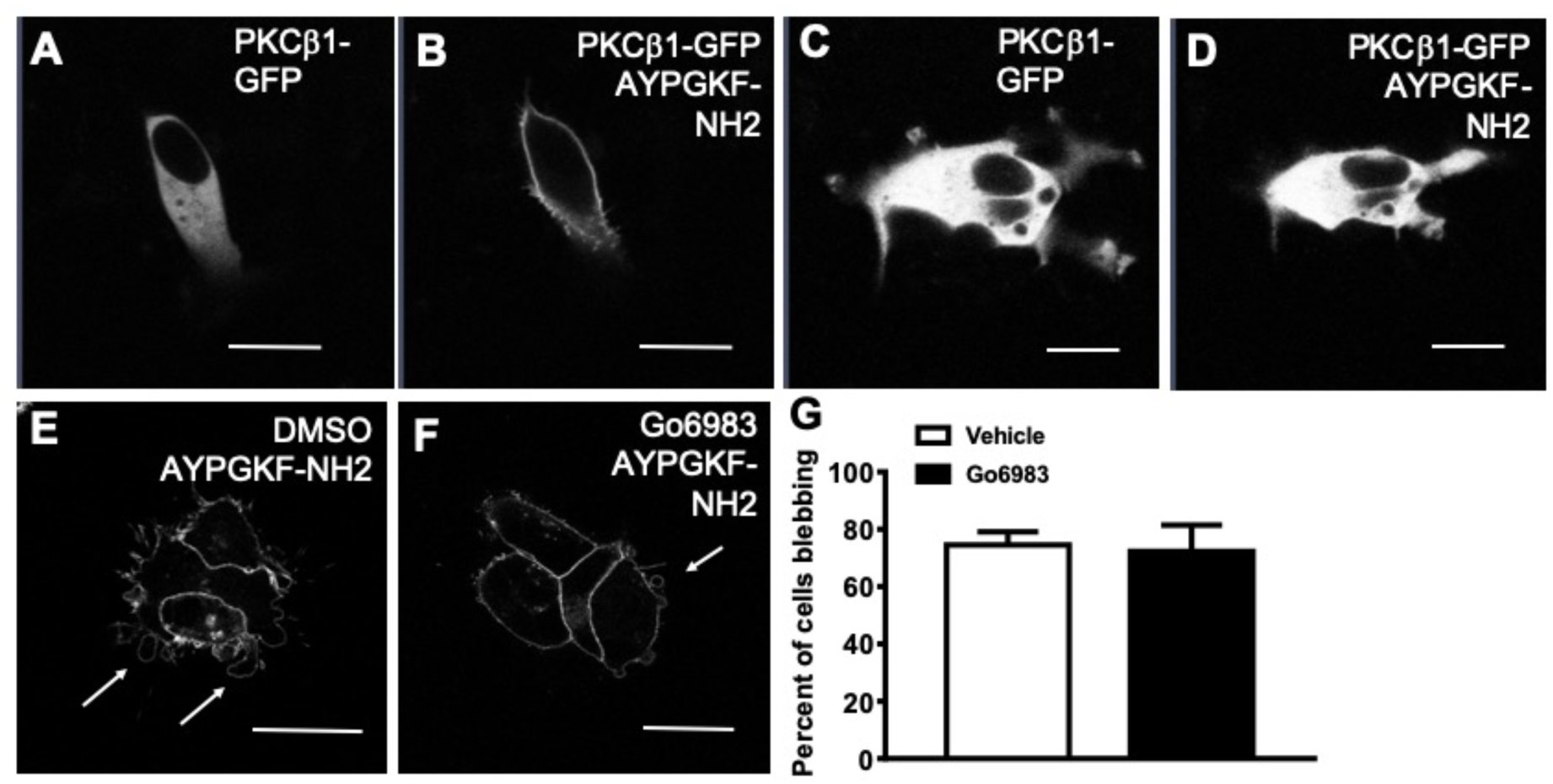
PAR4-mediated cell shape change is PKC-independent. HEK-293 cells transiently expressing either PAR4-mCherry (not shown) (A, B) or dRS-PAR4-mCherry (not shown) (C, D) with PKCβ1-GFP, were treated with 30 μM AYPGKF-NH_2_ and imaged by confocal microscopy, size bars are 20 μm. HEK-293 cells stably expressing PAR4-YFP were incubated with DMSO (E) or with 100 nM GO6983 for 15 minutes (F) prior to 2 a minute stimulation with 30 μM AYPGKF-NH_2_ and subsequent imaging by confocal microscopy. Cells were scored for blebbing or non-blebbing, graph shows mean +/- SEM, n = 4, not significantly different, t-test (G).

After ruling out Gα_q/11_ as a potential signalling partner for PAR4-mediated membrane blebs, we tested Gα_i_ recruitment as a potential regulator of these responses through inhibition of Gα_i_ signalling with pertussis toxin. HEK-293 cells stably expressing PAR4-YFP were incubated with pertussis toxin (100 nM) for 18 hrs prior to stimulating the cells with 30 μM AYPGKF-NH_2_. We did not observe any reduction in the number of cells that displayed blebbing when compared to cells incubated with vehicle control (saline), 76.3% +/-4.7 and 78.0% +/-3.4 respectively (Fig. 5A-C).

**Figure 5.**
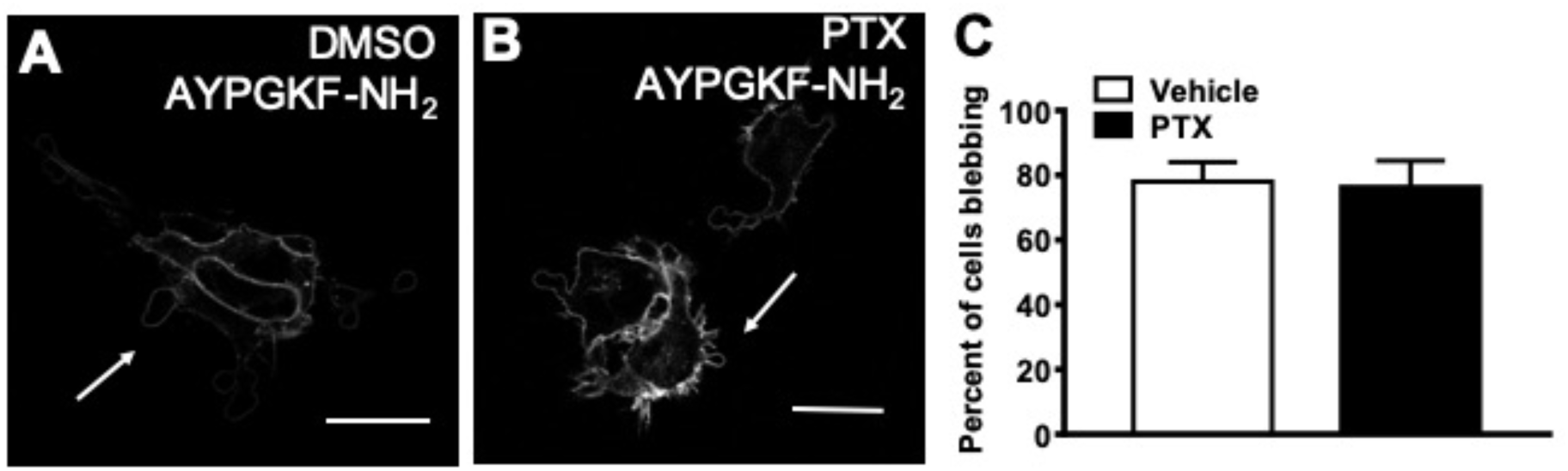
PAR4-mediated cell shape change is Gα_i_-independent. HEK-293 cells stably expressing PAR4-YFP were incubated in either DMSO (A) or 100 nM PTX (B) for 18 hours prior to 30 μM AYPGKF-NH_2_ treatment for 2 minutes and confocal imaging. Arrows show bleb formation, size bars are 20 μm. Graphs shows mean +/- SEM, n = 3, not significantly different, Mann-Whitney (C).

### PAR4-mediated cell shape change is RhoA and ROCK dependent

Since RhoA can be activated by a Gα_q/11_ and β-arrestin-1-dependent mechanism following activation of the angiotensin II type 1 receptor (Barnes *et al.*, 2005) and since RhoA is also well-established as a regulator of actin cytoskeleton rearrangements (Barnes *et al.*, 2005; Aoki *et al.*, 2016), we examined whether PAR4-mediated blebbing was RhoA- and ROCK-dependent. Treatment of PAR4-YFP expressing HEK-293 cells with the ROCK specific inhibitor GSK269962 (Stavenger *et al.*, 2007) significantly reduced the number of blebbing cells. 72.53% +/-6.24 of DMSO vehicle control treated cells displayed blebbing while only 15.6% +/-6.28 of cells treated with GSK269962 (100 nM) displayed bleb formation in response to 30 μM AYPGKF-NH_2_ (Fig. 6A-C). To further confirm the role of RhoA in PAR4-mediated changes in the plasma membrane, a RhoA knock out HEK-293 cell line (RhoA-KO HEK) was created by use of CRISPR/Cas9 targeting. RhoA-KO HEK cells and control HEK-293 cells were transiently transfected with PAR4-YFP, stimulated with 30 μM AYPGKF-NH_2_ and imaged by confocal microscopy (Fig. 6D-F). 75% +/-7.8 of control HEK-293 cells expressing PAR4-YFP and treated with AYPGKF-NH_2_ displayed membrane blebbing while RhoA-KO HEK expressing PAR4-YFP show significantly fewer cells with blebs in response to agonist (22.8% +/-6.2) (Fig. 6F). Taken together, these data indicate that stimulation of PAR4-YFP elicits cell shape changes that are blocked by blebbistatin and by the ROCK inhibitor GSK269962. Further, these cell shape changes do not occur in cells that do not express RhoA protein, suggesting that this signalling pathway is a RhoA and ROCK dependent pathway.

**Figure 6.**
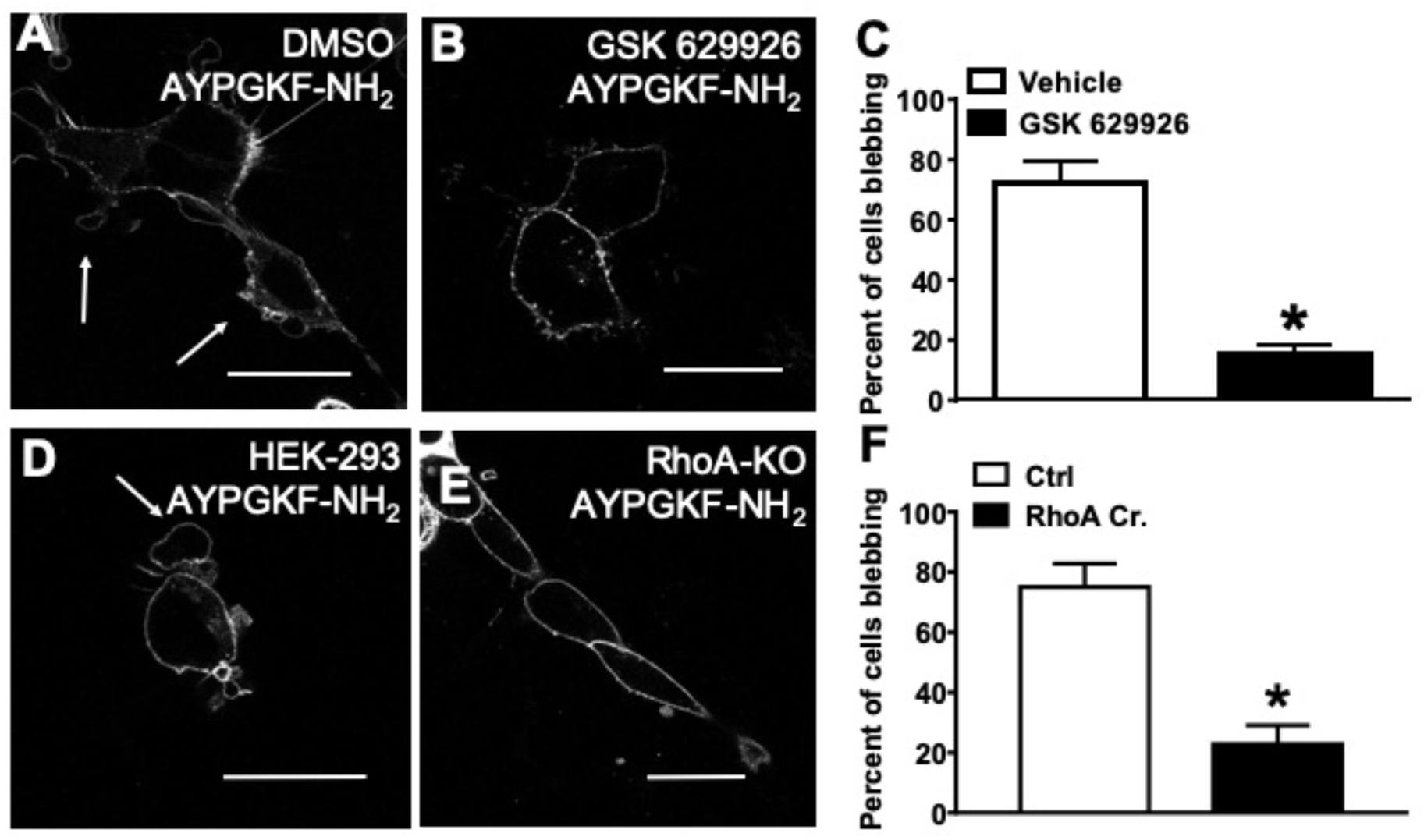
PAR4-mediated cell shape change is RhoA- and ROCK-dependent. HEK-293 cells stably expressing PAR4-YFP were incubated in either DMSO (A) or 100 nM GSK269962 for 1 hour (B) prior to a 2 minute treatment with 30 μM AYPGKF-NH_2_ followed by confocal microscopy. Arrows show bleb formation, size bars are 20 μm. (C) graph shows mean +/- SEM, n = 4, asterisk shows significantly different, Mann-Whitney, p<0.05. HEK-293 cells and RhoA-KO HEK transiently expressing PAR4-YFP were stimulated with 30 μM AYPGKF-NH_2_ for 2 minutes prior to confocal imaging (D, E). (F) Graph shows mean +/- SEM, n = 4, asterisk shows significantly different, Mann-Whitney, p<0.05.

To confirm that these cellular responses can also be recapitulated by a physiological PAR4 agonist, we tested the ability of thrombin to elicit similar cell shape changes. Since thrombin can also activate PAR1 which is endogenously expressed in HEK-293 cells we conducted these experiments in PAR1 knockout HEK-293 cells (PAR1-KO HEK) (Mihara *et al.*, 2016) stably expressing PAR4-YFP (PAR1-KO-HEK-PAR4-YFP). PAR1-KO-HEK-PAR4-YFP cells were treated with 3 U/ml thrombin and visualized by confocal microscopy. Treatment of cells with thrombin caused cell shape changes similar to what was observed in HEK-293-PAR4-YFP cells treated with AYPGKF-NH_2_ (Fig. 7A). PAR1-KO-HEK-PAR4-YFP cells were then treated with blebbistatin prior to thrombin stimulation (Fig. 7B). Blebbistatin significantly reduced the number of PAR1-KO-HEK-PAR4-YFP cells displaying membrane blebs from 68.3% +/-4.6 to 11.67% +/-5.78 (Fig. 7D). Finally, PAR1-KO-HEK-PAR4-YFP cells were treated with GSK269962 prior to stimulation with thrombin (Fig. 7C). GSK269962 (100 nM) significantly reduced the number of cells that displayed blebbing in response to thrombin stimulation (Fig. 7D). Together these data indicates that cell blebbing is triggered not only by the synthetic PAR4 activating peptide AYPGKF-NH_2_, but also by one of the endogenous activators of PAR4, thrombin, in a PAR1 null background. Cell shape changes mediated by thrombin activation of PAR4 are also RhoA- and ROCK-dependent.

**Figure 7.**
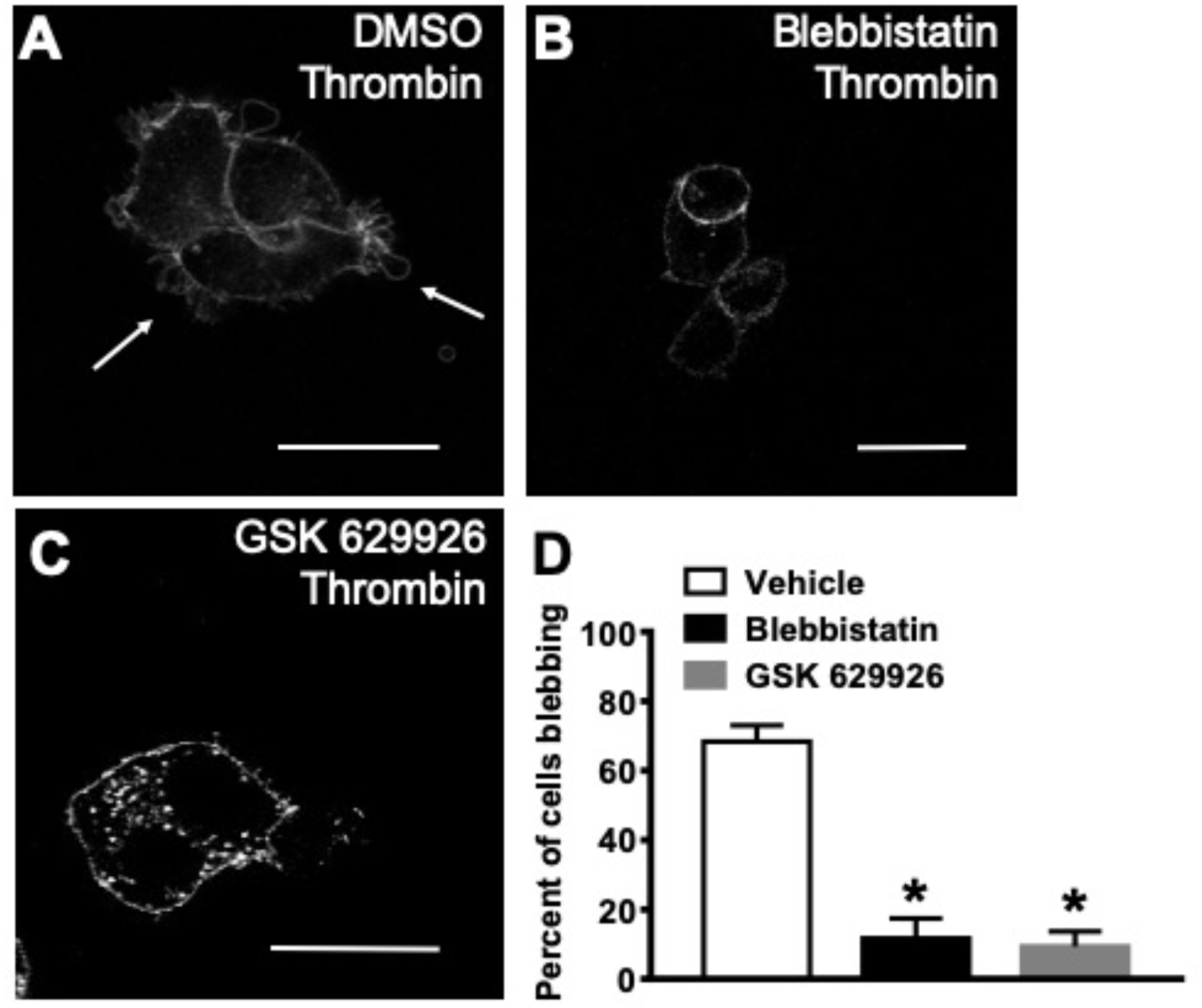
Thrombin-induced PAR4-mediated cell shape change is ROCK-dependent. PAR1-KO-PAR4-YFP-HEK-293 cells were incubated in either DMSO (A) or 10 μM blebbistatin for 15 minutes (B) or 100 nM GSK269962 for 1 hour (C) prior to a 2 minute treatment with 30 μM AYPGKF-NH_2_ followed by confocal microscopy. Arrows show bleb formation, size bars are 20 μm. Graph shows mean +/- SEM, n = 4, asterisk shows significantly different, Mann-Whitney, p<0.05 (D).

### PAR4-mediated cell shape change is β-arrestin dependent

Since the non-blebbing dRS-PAR4-YFP expressing cells are also deficient in β-arrestin recruitment (Ramachandran *et al.*, 2017), we next examined the contribution of β-arrestin-mediated signalling in PAR4-dependent cell membrane blebbing. To elucidate a role for β-arrestins in PAR4-mediated cell shape change, a β-arrestin-1 and -2 double knockout cell line (β-arrestin-1/-2 KO HEK) was created using CRISPR/Cas9 targeting. β-arrestin-1/-2 KO HEK cells transiently expressing PAR4-YFP (β-arrestin-1/-2 KO HEK-PAR4-YFP) were treated with 30 μM AYPGKF-NH_2_ and visualized by confocal microscopy. A significant reduction in the number of blebbing β-arrestin-1/-2 KO HEK-PAR4-YFP (46%+/-8.2) was observed when compared to control HEK-293 cells transiently expressing PAR4-YFP (80.8%+/-6.1)(Fig. 8A-C).

**Figure 8.**
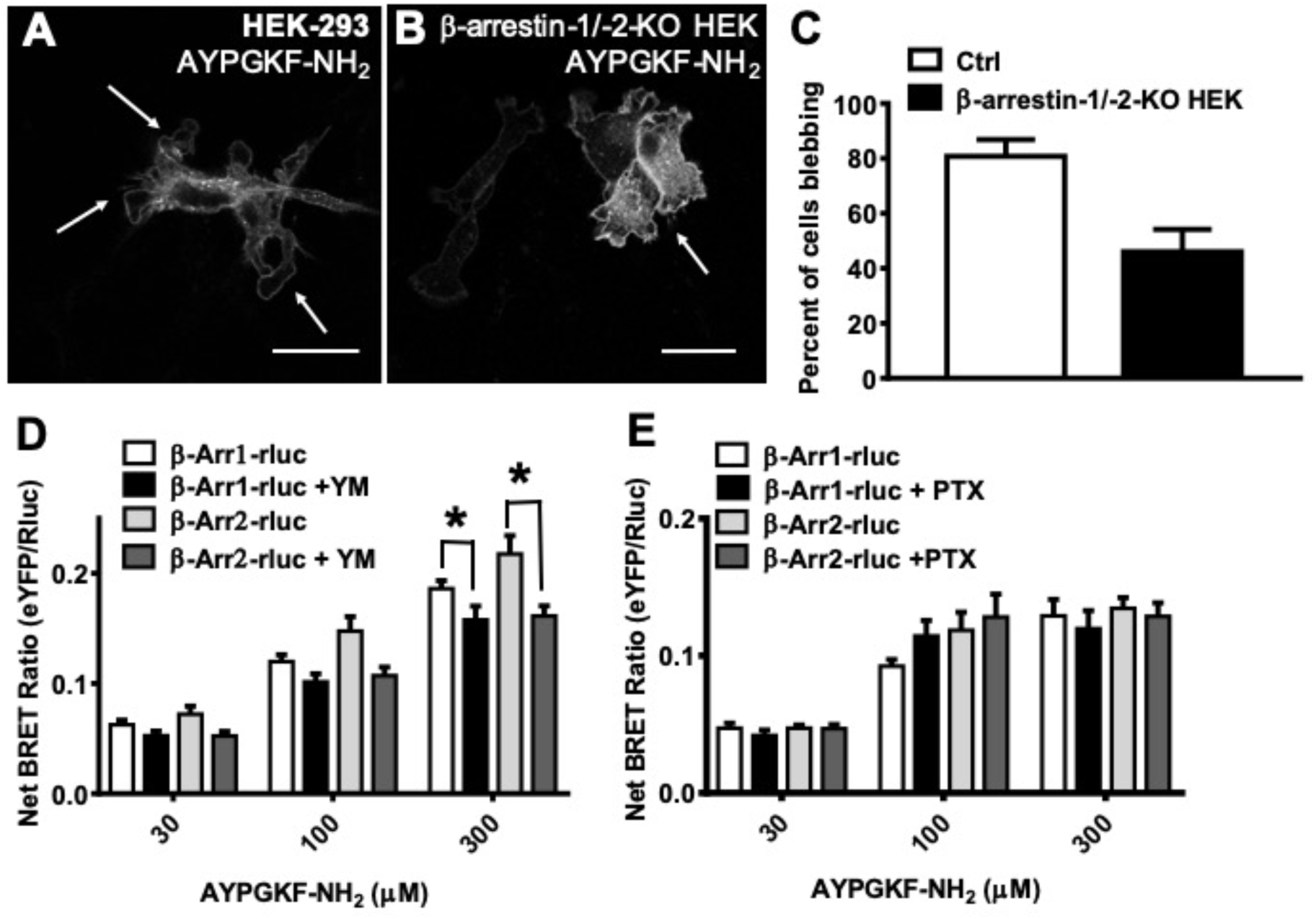
PAR4-mediated cell shape change is β-arrestin-dependent. HEK-293 cells (A) or β-arrestin-1/-2-KO HEK transiently expressing PAR4-YFP (B) were stimulated in 30 μM AYPGKF-NH_2_ for 2 minutes prior to confocal imaging. Arrows show bleb formation, size bars are 20 μm. (C) Graph shows mean +/- SEM, n=3, asterisk shows significantly different, t-test, P<0.05). HEK-293 cells transiently expressing PAR4-YFP with either β-arrestin-1-Rluc or -2-Rluc were incubated in DMSO or 100 nM YM254890 (D) or 100 nM PTX for 18 hours (E) for 20 minutes prior to testing in the BRET assay. Graph shows mean +/- SEM, n = 4, asterisk shows significantly different, 2-way ANOVA.

Since recent reports have questioned whether β-arrestin mediated signalling can occur in the absence of G-protein activation (Grundmann *et al.*, 2018) we sought to understand this requirement in the context of PAR4 signalling. Even though Gα_q/11_ and Gα_i_ recruitment was not implicated in PAR4 dependent membrane blebbing, we nevertheless examined the effect of blocking these pathways on β-arrestin-1/-2 recruitment to PAR4. To this end we blocked Gα_q/11_ with the inhibitor YM-254890 (100 nM) and blocked Gα_i_ with pertussis toxin (100 nM) and examined their ability to disrupt β-arrestin recruitment to PAR4-YFP. We employed a BRET assay to monitor interaction between PAR4-YFP and β-arrestin-1-Rluc or β-arrestin 2-Rluc in response to 30, 100 and 300 μM AYPGKF-NH_2_. Treatment of cells with YM-254890 did not significantly reduce recruitment of β-arrestin-1 or β-arrestin-2 recruitment to PAR4 at 30 μM or 100 μM concentrations of AYPGKF-NH_2_ but did significanctly reduce recruitment of both β-arrestin-1 and -2 at the 300 μM AYPGKF-NH_2_ (Fig. 8D) treatment condition. Treatment of cells with pertussis toxin had no effect on β-arrestin recruitment at any of the concentration tested (Fig. 8E).

Finally, we tested a role for Gβγ signalling in PAR4-mediated cell shape change using the Gβγ inhibitor gallein (10 µM). HEK-293 cells stably expressing PAR4-YFP were incubated with gallein or DMSO vehicle control prior to treatment with 30 μM AYPGKF-NH_2_. Cells treated with gallein did not show a significant reduction in membrane blebbing compared to DMSO treated cells, 61.75% +/-4.5 and 85.28% +/-5.2 (Fig. 9A-C). Cells expressing PAR4-YFP and either β-arrestin-1-Rluc or -2-Rluc incubated with gallein prior to treatment with AYPGKF-NH_2_ at 30, 100 and 300 μM also retained their ability to recruit β-arrestin-1 and -2 at all concentrations of AYPGKF-NH_2_ (Fig. 9D). Taken together these data show that PAR4-mediated cell shape changes is independent of Gα_q/11_, Gα_i_ and Gβγ. Further inhibition of Gα_i_ and Gβγ individually does not impede β-arrestin recruitment to PAR4, while Gα_q/11_ inhibition partially reduces β-arrestin recruitment to PAR4.

**Figure 9.**
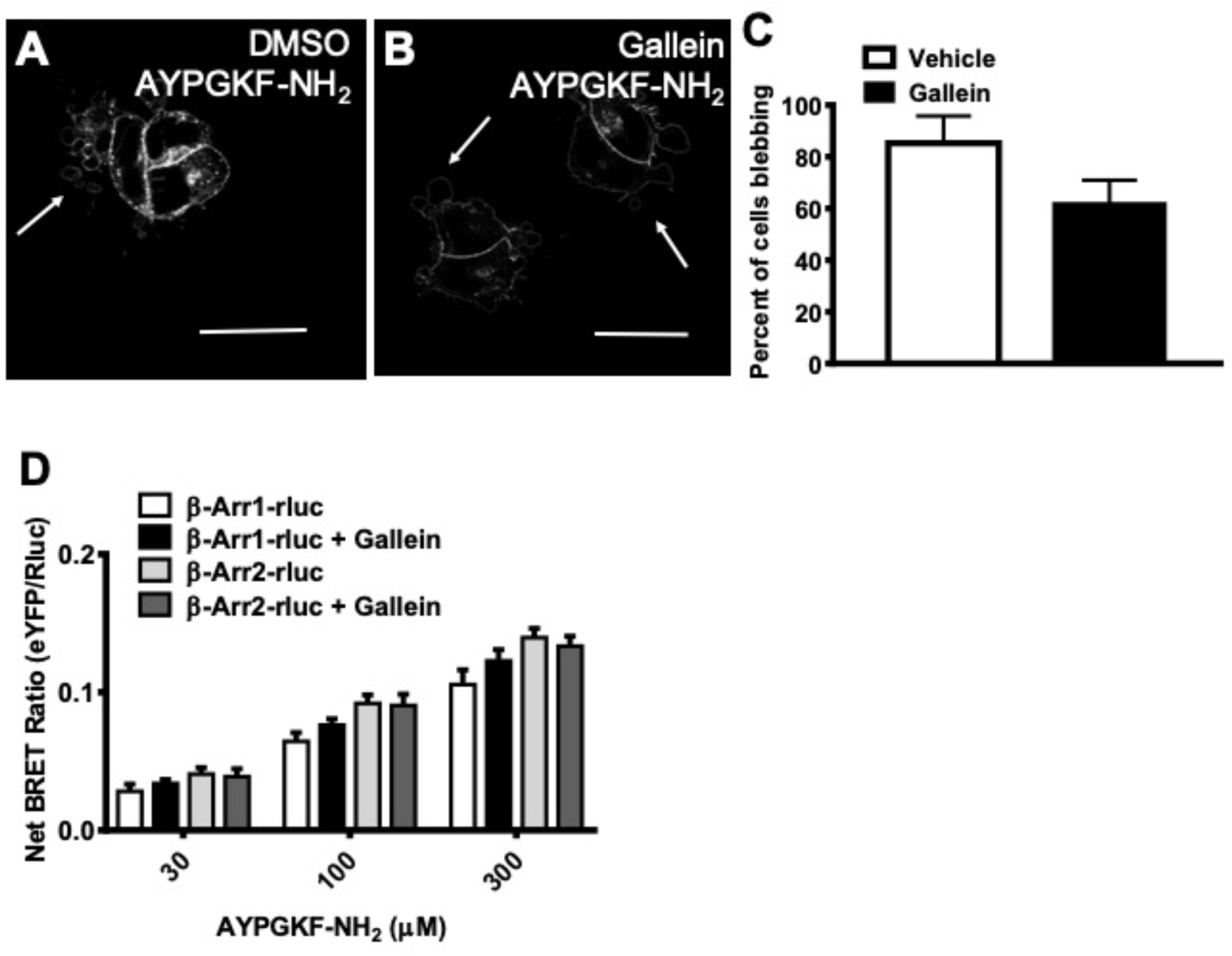
PAR4-mediated cell shape change is Gβγ-independent. HEK-293 cells stably expressing PAR4-YFP were incubated in either DMSO (A) or 10 μM gallein for 20 minutes prior to 30 μM AYPGKF-NH_2_ for 2 minutes and confocal imaging. Arrows show bleb formation, size bars are 20 μm. Graphs shows mean +/- SEM, n=4, not significantly different, Mann-Whitney (C). HEK-293 cells transiently expressing PAR4-YFP with either β-arrestin 1-Rluc or -2-Rluc were incubated in DMSO or with 10 μM gallein for 20 minutes (H) prior to testing in the BRET assay. Graph shows, mean +/- SEM, n = 4, not significantly different 2-way ANOVA.

### PAR4 activation in rat primary aortic vascular smooth muscle cells leads to cell shape changes

In order to examine whether PAR4 activation triggered cell blebbing in cells that endogenously express PAR4, we turned to the rat vascular smooth muscle cells. Vascular smooth muscle cells are reported to express PAR4 (Bretschneider *et al.*, 2001; Dangwal *et al.*, 2011) and in our hands rat aortic smooth muscle cells from both WKY and SHR rats expressed PAR4 (Fig. 10I). We labelled the smooth muscle cell membrane with cell mask (ThermoFisher) and stimulated with 30 μM AYPGKF-NH_2_. Both WKY and SHR VSMC displayed cell shape changes resembling membrane blebs in response to PAR4 agonist treatment (Fig 10A, E). In order to establish whether the signalling mechanism for these cell shape changes was the same in both VSMC and HEK-293 cells, we pretreated VSMC with blebbistatin and the ROCK inhibitor GSK269962 prior to activating PAR4 with 30 μM AYPGKF-NH_2_. 67.25 +/-6.6% of WKY cells display cell shape changes in response to PAR4 agonist, (Fig. 10A), which was significantly reduced with treatment blebbistatin (9.0 +/-3.1%) and with treatment with GSK269962 (6.5 +/-0.28%) (Fig. 10B-D). Consistent with these findings, 65.75 +/-5.4% of SHR cells also displayed blebbing following PAR4 activation and these responses were significantly reduced with blebbistatin (14.5 +/-1.9%) and with GSK269962, (15.5 +/-3.1%) treatment (Fig. 10F-H). These data indicate that endogenous PAR4 activation elicits membrane blebbing in VSMC that is ROCK dependent in keeping with our findings in the HEK-293 cells exogenously expressing PAR4.

**Figure 10.**
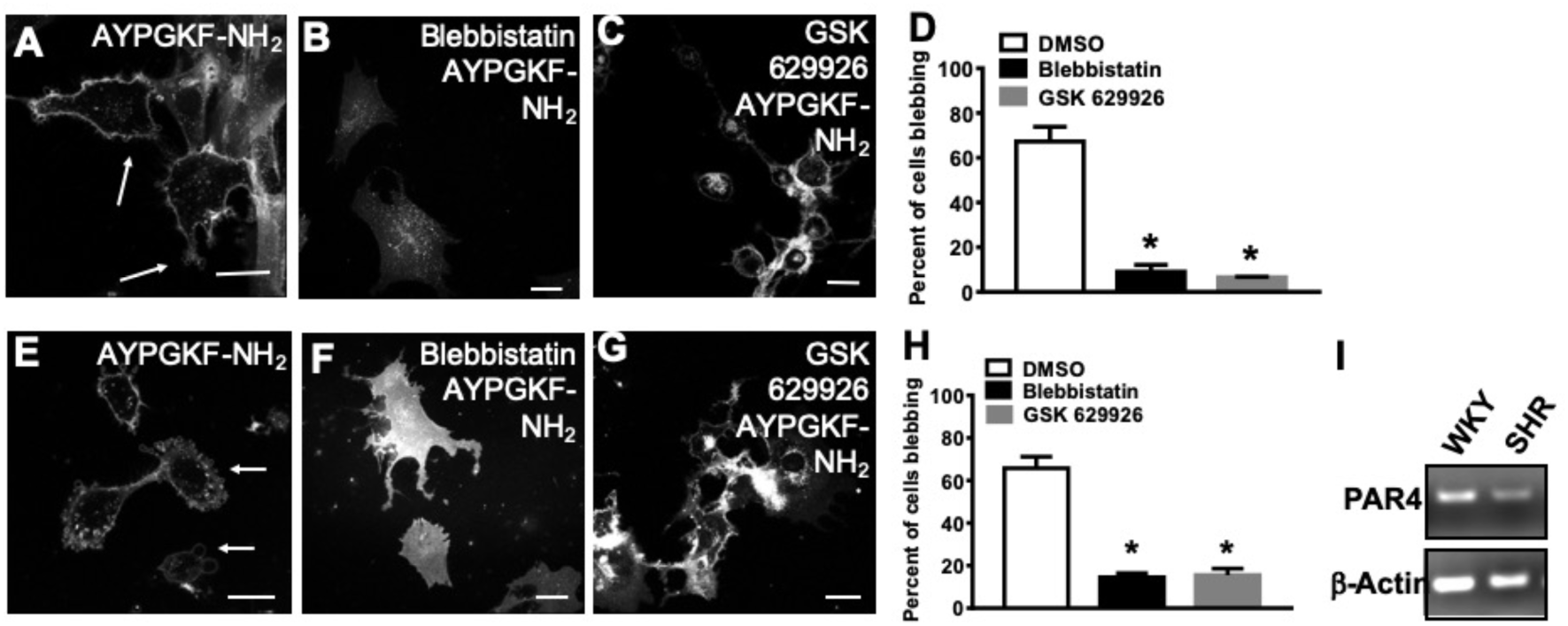
PAR4 mediates membrane blebs in vascular smooth muscle cells. Primary cultured vascular smooth muscle cells derived from Wistar Kyoto (WKY) (A-D) or from spontaneously hypertensive (SHR) (E-H) rats were pre-incubated for 10 minutes in cell mask to stain the plasma membrane, cells were then treated with 30 μM AYPGFK-NH_2_ and imaged by confocal microscopy. WKY cells and SHR cells were preincubated with blebbistatin or with GSK269962 prior to stimulation with AYPGFK-NH_2_ (B, C, F, G), arrows show bleb formation, size bars are 20 μm. Images were scored for blebbing or non-blebbing (D and H), graph shows mean +/- SEM, n = 4, asterisk shows significantly different, ANOVA, p<0.001.

### PAR4 mediated β-arrestin recruitment influences gene transcription

β-arrestins are scaffold proteins that couple GPCRs to multiple signalling cascades in a manner that is differential to heterotrimeric G-protein coupling (Laporte and Scott, 2019). Since PAR4-mediated membrane blebbing is β-arrestin-dependent, we sought to take a complementary approach to explore PAR4-mediated β-arrestin-dependent pathways. In order to do this, we compared the transcriptomes of HEK-293 and β-arrestin-1/-2-KO HEK cells activated with 30 μM AYPGKF-NH_2_ for 3 hours as well as the transcriptomes of untreated HEK-293 and β-arrestin-1/-2-KO HEK cells transiently expressing PAR4-YFP. Principal component analysis and hierarchical clustering of gene expression profiles shows clear clustering of HEK-293 samples as well as treated and untreated β-arrestin-1/-2-KO HEK samples (Fig. S1A-D). Differential expression analysis between the HEK-293 cells and β-arrestin-1/-2-KO HEK cells yielded 2,764 and 3,155 significantly differentially expressed genes (P_adj_ <0.05 and Fold-Change>2) between the HEK-293 and β-arrestin-1/-2-KO HEK samples in untreated and treated conditions, respectively. A total of 1,313 differentially expressed genes were shared between the untreated and treated conditions (Fig. 11). Of note, even though we see a depletion of β-arrestin protein in our β-arrestin-1/-2-KO HEK cell line, we do not see down regulation of β-arrestin-1 or -2 transcripts in the RNA seq profile. Gene ontology (GO) enrichment analysis of the significantly upregulated genes in samples treated with 30 μM AYPGKF-NH_2_ (n = 1506) revealed biological process terms related to cell-cell adhesion, and G-protein coupled receptor signalling pathway which are consistent with our finding of membrane blebbing triggered by PAR4 activation. (Table 1). Additionally, GO terms relating to known functions of PAR4 including neuron-neuron synaptic transmission, blood coagulation, blood circulation, and sensory perception were also found to be enriched in upregulated genes (Table 1). GO analysis of the downregulated genes (Treated only: n = 336) revealed biological process terms related to intracellular signal transduction. (Table 2). Taken together, these results suggest that β-arrestin-1/-2 are important regulators of signalling and gene transcription following PAR4 activation. Genes upregulated relating to cell-cell adhesion and GPCR signalling are consistent with our finding of β-arrestin-dependent membrane blebbing following PAR4 activation.

## Discussion

We have demonstrated that activation of PAR4 with thrombin or the synthetic PAR4 activating peptide, AYPGKF-NH_2_, causes a rapid cell shape change response in PAR4-transfected HEK-293 cells or in vascular smooth muscle cells that endogenously express PAR4. Cell shape changes are pharmacologically inhibited by blebbistatin and are consistent with membrane bleb formation. We observed membrane blebs that formed within 2-5 minutes of agonist treatment and lasted for upto 30 minutes. Membrane blebbing could be pharmacologically inhibited by the ROCK inhibitor GSK269962 or through CRISPR/Cas9 mediated knockout of RhoA. CRISPR/Cas9 knockout of β-arrestin-1 and -2 partially abolished PAR4-dependent mebrane blebbing. We further found PAR4-dependent membrane bleb formation to be independent of Gα_q/11_, Gα_i,_ Gβγ, and PKC. Overall, our data suggest that RhoA-dependent membrane blebbing occurs downstream of PAR4 activation and β-arrestin recruitment.

Non-apoptotic cell membrane blebbing plays an important role in various physiological and pathologlical processes. Various stimuli have been reported to trigger bleb formation leading to cellular responses including enhanced cell motility, invasion, cell locomotion, and regulation of cell polarity in embryonic development (Charras and Paluch, 2008; Fackler and Grosse, 2008; Ikenouchi and Aoki, 2016). Blebbing is also an important regulator of wound healing, immune cell maturation, and inflammation. In this context, the PAR family of GPCRs is well established as critical regulators of the innate immune response to injury and infection. PARs also elicit cellular responses that allow coagulation cascade enzymes such as thrombin and other serine proteinases to regulate various cellular functions. PAR1 and PAR4 serve as the receptors for thrombin on human platelets, though these receptors regulate different aspects of platelet activation (Coughlin, 1999; Kahn *et al.*, 1999; Ma *et al.*, 2005; Holinstat *et al.*, 2006; Voss *et al.*, 2007). PAR4 is described as the low-affinity thrombin receptor on human platelets. This lower affinity stems from the lack of a hirudin like binding site for thrombin on PAR4, that is present on PAR1. In platelets, PAR4 activation typically requires a much higher concentration of thrombin to be present and PAR4 activation typically results in more sustained calcium signalling compared to PAR1-dependent signalling (Covic *et al.*, 2000; Shapiro *et al.*, 2000).

Both PAR1 and PAR4 activation triggers platelet aggregation, with PAR4 signalling critical for full platelet spreading and formation of stable aggregates. In rodents, PAR4 serves as the sole thrombin receptor in platelets (Sambrano *et al.*, 2001). It has been previously shown in Gα_q/11_ knock out mice, that a thrombin activated Gα_q/11-_mediated scalcium response is necessary for platelet aggregation (Offermanns *et al.*, 1997). However, platelets from the mouse Gα_q/11_ knockout retained the ability to change shape in response to thrombin stimulation. Similar findings were reported in platelets treated with a small molecule Gα_q/11_ antagonist UBO-QIC showing that platelet aggregation was inhibited without affecting shape change responses (Inamdar *et al.*, 2015). Consistent with these findings, we have observed a cell shape change specific to PAR4 activation that is G_q/11-_ independent and a RhoA-mediated phenomenon. Recent evidence suggests that PAR4 plays an important role in promoting platelet granule release and platelet-leukocyte interactions (Rigg *et al.*, 2019), responses which also rely on a RhoA-mediated cytoskeletal rearrangement (Moers *et al.*, 2003; Aslan and Mccarty, 2013). These findings raise the interesting possibility that different signalling pathways may underlie PAR4 regulation of distinct aspects of platelet activation and further study is required to fully elucidate the role of different signalling cascades in mediating PAR4 responses in platelets. Our studies suggest that in HEK cells the Gα_q/11_-coupled pathway and the RhoA pathway, which in this instance is likely downstream of Gα_12/13_ coupling, can act independently and therefore may be independent targets for pharmacological manipulation.

PAR4 expression has also been reported in other cell types involved in the response to injury including endothelial cells and smooth muscle cells (Bretschneider *et al.*, 2001; Hamilton *et al.*, 2001; Fujiwara *et al.*, 2005; Ritchie *et al.*, 2007). Here we demonstrate that agonist stimulation of endogenous PAR4 in vascular smooth muscle cells leads to membrane blebbing. This finding is consistent with a previous study which showed that another GPCR AT1R mediates membrane blebbing by a RhoA-dependent mechanism in a vascular smooth muscle cell line (Godin and Ferguson, 2010). RhoA is well established as a mediator of changes in the plasma membrane actin cytoskeleton and in regulating GPCR mediated cell shape change (Barnes *et al.*, 2005; Godin and Ferguson, 2010; Aoki *et al.*, 2016). Our findings add PAR4-dependent signalling to the cell surface signalling molecules that can trigger this pathway.

β-arrestins are now well established as important molecular scaffolds linking GPCRs to not only molecular endocytic partners to facilitate receptor endocytosis, but also to second messenger signal cascades (Ferguson, 2001; Magalhaes *et al.*, 2012). For example β-arrestins link GPCRs to p44/42 MAP kinase signalling from the endosome (Luttrell *et al.*, 2001; Luttrell and Lefkowitz, 2002). PAR4 couples to both β-arrestin-1 and β-arrestin-2 and PAR4-mediated phosphorylation of AKT in platelets is β-arrestin-2-dependent (Li *et al.*, 2011). We explored β-arrestin-1/-2-dependent PAR4 signalling by using RNA sequencing analysis in HEK cells and β-arrestin knock out HEK cells treated or not with the PAR4 specific agonist peptide AYPGKF-NH_2_. Our data shows that β-arrestins can regulate PAR4-dependent gene transcription. Consistent with our finding that PAR4 mediates membrane bleb formation, we see an upregulation of gene ontology terms in cell-cell adhesion, blood coagulation and blood circulation. It is well established that PAR4 activation regulates platelet response, thus links to gene ontology terms related to blood coagulation are perhaps unsurprising and uncover an important link to β-arrestin in these processes. Our RNA-seq analysis uncovered a number of other additional signalling pathways that could be linked to PAR4 and β-arrestin such as neuron-neuron synaptic transmission (GO:0007270), natural killer cell activation (GO:0030101) and sensory perception (GO:0007600). PAR4 expression has been reported in various neuronal and immune cells and these data provide impetus to further examine the roles of PAR4 in these cells (Henrich-Noack *et al.*, 2006; Russell *et al.*, 2009; Peng *et al.*, 2019). More broadly were also interested in understanding if PAR4 signalling to β-arrestin could proceed independently of G-protein coupling. Blocking Gα_q/11_ partially inhibited β-arrestin recruitment to PAR4 while blocking of Gα_i_- or βγ-mediated signalling had no effect. It however remains to be determined whether β-arrestin recruitment dependent signalling downstream of PAR4 is truly independent of G-protein coupling or occurs following Gα_12/13_ activation.

In conclusion, we have uncovered a PAR4-mediated cellular response that is independent of Gα_q/11_ coupling and occurs downstream of RhoA activation and β-arrestin signalling. These data provide further evidence for pathway-selective signalling responses through PAR4 and may guide future development of PAR4 targeting strategies.

## Figure Legends

**Supplementary Figure 1.**
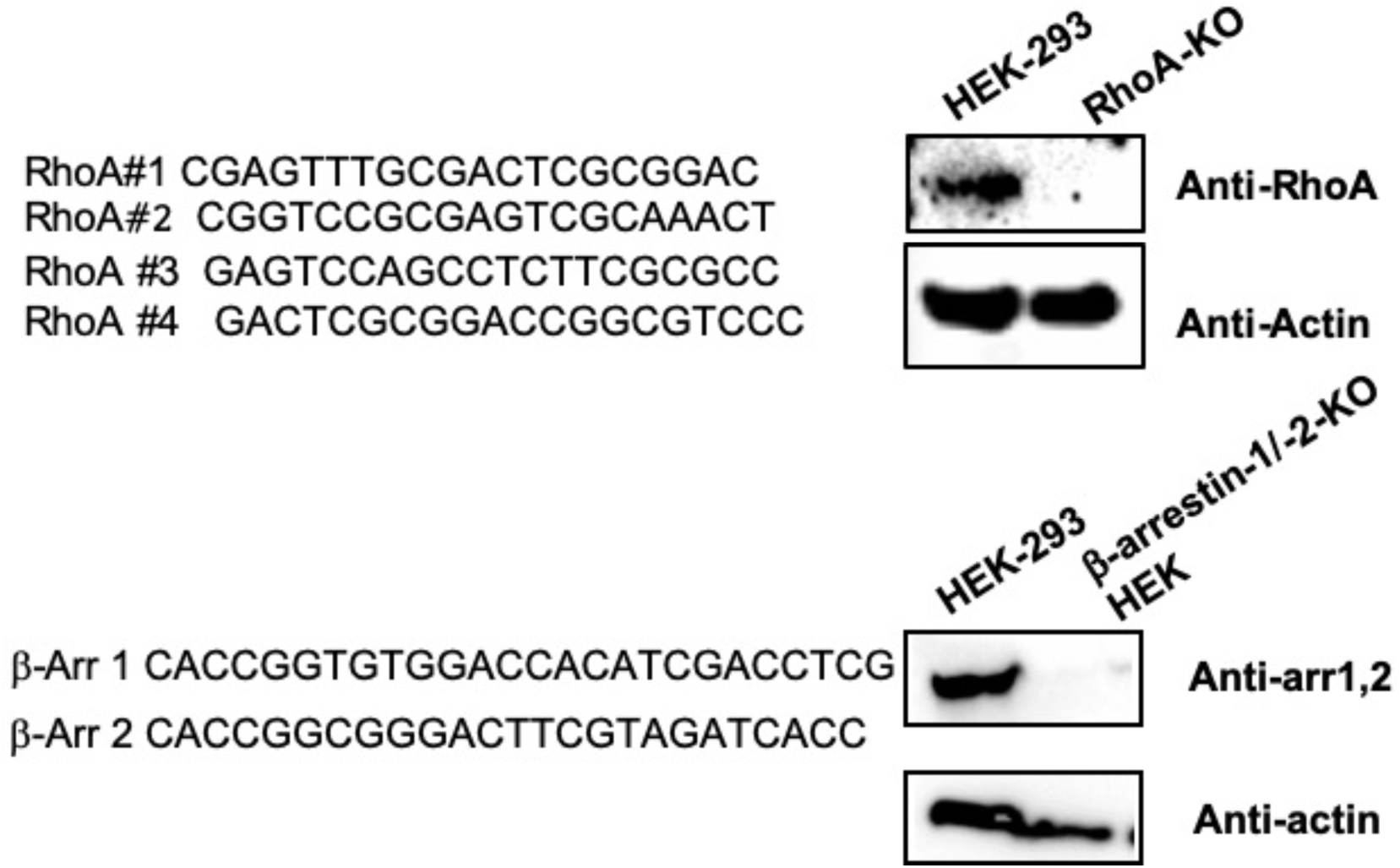
CRISPR/Cas9 knock out of β-arrestin-1/-2 and RhoA in HEK-293 cells. HEK-293 cells were transiently transfected with either guide RNA specific to β-arrestin-1 and β-arrestin-2, or four guide RNAs specific to RhoA. Cells expressing pSpCas9(BB)-2A-GFP encoding guide RNAs were selected for using FACS Aria cell sorter and subsequently knockout was determined by western blot.

**Supplementary Figure 2.**
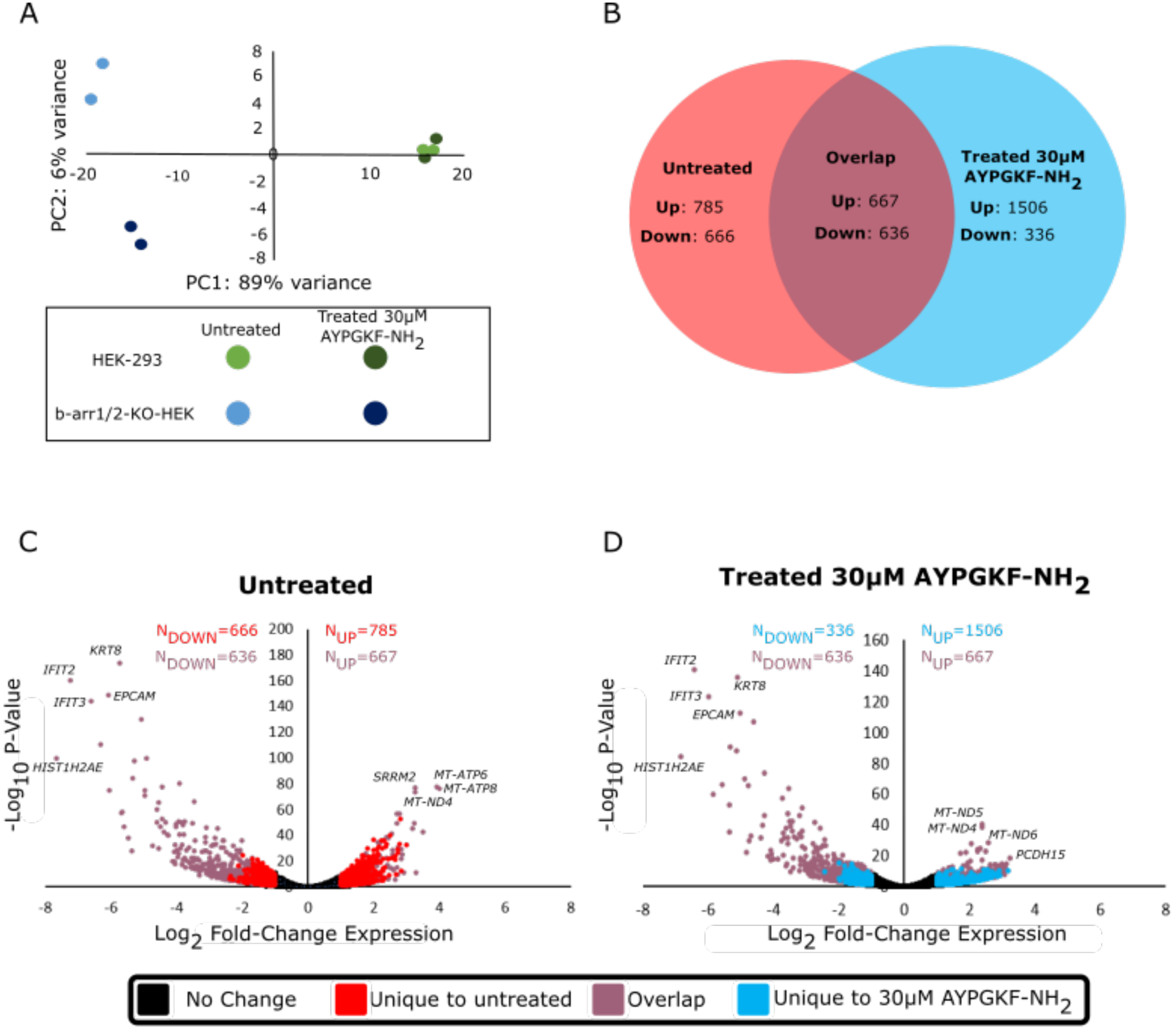
Knockout of β-arrestin-1/-2 results in differentially expressed genes. **(A)**Principal component analysis biplot showing the first two principal components (PCs) that account for most of the variance between samples. **(B)** Venn diagram showing overlap of differentially expressed genes (P_adj_<0.05 and Fold-Change>2) in β-arrestin-1/-2-KO HEK cells treated with PAR4 agonist (Treated) or with vehicle (Untreated) when compared to similarly treated HEK-293 cells. **(C & D)** Volcano plots showing significantly differentially expressed genes between β-arrestin-1/-2-KO and HEK-293 cells when treated with vehicle **(C)** or 30 μM AYPGKF-NH_2_ **(D)**. Of the differentially expressed genes, 666 were downregulated and unique to untreated cells, 336 were downregulated and unique to treated cells, and 636 were found to be downregulated in both conditions. Additionally, 785 genes were upregulated and unique to untreated cells, 1506 were upregulated and unique to treated cells, and 667 were found to be upregulated in both conditions.

